# Sensory recurrence and convergent integration in the *C. elegans* connectome

**DOI:** 10.64898/2026.10.29.762790

**Authors:** Elkaïoum M. Moutuou, Habib Benali

## Abstract

The synaptic wiring of *Caenorhabditis elegans* has a global feed-forward architecture, from sensory to motor neurons. Yet, whole-brain optogenetic mapping shows that signals do not flow predominantly forward, and sensory neurons respond to more neurons than they drive. Where the wiring contains pathways leading back to their starting point is unclear. Here, we locate this recurrence using the feedback coefficient, derived from the Kubo–Martin– Schwinger framework for directed networks, which quantifies the proportion of walks reaching a neuron that originate from itself. The connectome has about three times more recurrence than strength-matched random networks, with the largest excess in sensory neurons, whereas random wiring places it on highly connected neurons. Along the thermotaxis pathway, this produces a recurrence gradient from thermosensory neurons to integrating interneurons. The ordering replicates in five reconstructions, including two independent adults, persists when reciprocal contacts are preserved or gap junctions removed, but vanishes with only short paths or loop counts. Convergence shows the reverse pattern, with every neuron receiving walks from fewer sources than in random wiring, especially sensory neurons. The wiring extends from a sensory periphery folding back on itself to interneurons and motor neurons integrating many sources, a candidate structural substrate for non-feed-forward signalling.

## Introduction

Complete reconstructions of the *C. elegans* nervous system in several animals [1, 2, 3, 4] have shown that the global structure of its synaptic connectivity is feed-forward; *i.e*., synapses lead on balance from sensory neurons through interneurons to motor neurons [5]. This holds, however, only for the net direction of the synapses since many connections are reciprocal and the most highly connected interneurons are densely linked to one another [6]. How signals propagate through this architecture had not been measured systematically until recently. Randi *et al*. [7] optogenetically stimulated single neurons while recording calcium activity across the head, and measured signal propagation between more than 23,000 neuron pairs. They found that signal propagation was poorly predicted by models based on structural connectivity, in part because many functionally connected pairs had no direct synaptic contact and part of the signalling depended on extrasynaptic transmission through dense-core vesicles. Comparing this functional atlas with the connectome, Dvali *et al*. [5] then found that the two networks share few communities and hubs, that functional connectivity, unlike the connectome, is not predominantly feed-forward, and that amphid sensory neurons have more incoming than outgoing functional connections.

These observations are difficult to reconcile with a feed-forward view of the wiring, and while extrasynaptic signalling accounts for part of the discrepancy, the synaptic network itself may also contribute to it. Indeed, reciprocal and other recurrent connections, through which the activity of a neuron can return to it via other neurons, occur more often than expected by chance in cortical circuits [8] and in the *C. elegans* connectome [9], and in cortex they support persistent activity and the amplification and selection of sensory input [10, 11, 12]. If the *C. elegans* wiring contained many pathways leading back to their source neuron, a net feed-forward summary would conceal them, and signals could return towards the sensory periphery through synapses alone. Where such recurrent pathways are concentrated along the sensory-to-motor organisation of the network, and whether their distribution follows simply from the degree of each neuron, has not been established. This question, however, cannot be answered by simply counting loops, because the number of closed walks through a neuron grows with its number of connections, so that highly connected neurons accumulate the most loops even in random networks. A suitable measure should therefore compare the walks that return to a neuron with all the walks that reach it, and should be tested against networks in which every neuron keeps its number of connections.

The Kubo–Martin–Schwinger (KMS) framework for directed networks [13, 14] provides such a measure. It is built on the resolvent *R*_*β*_ = (*I − e*^*−β*^ *A*)^*−*1^, the same matrix that underlies Katz centrality [15], but uses it in two different ways. Its columns and rows are normalised into probability distributions over target and source neurons, which in the KMS construction are the pure states of the graph algebra of the directed multigraph underlying the connectome [13], whereas Katz centrality sums the resolvent into a single score per neuron. The inverse temperature *β* is also measured relative to a critical value *β*_*c*_ = log *r*(*A*), set by the spectral radius *r*(*A*) of the network, at which the walk sum diverges, so that the attenuation is defined by the network itself and is not a free parameter. The receptance profile of a neuron is the distribution, over source neurons, of the walk mass reaching it through walks of any positive length. Its self-entry, which we call the feedback coefficient **fb**, is the share of this incoming walk mass generated by the neuron itself. Specifically, **fb** measures recurrence relative to convergence, which loop counts alone do not capture.

We used **fb** to map recurrence across the whole somatic connectome and, in more detail, along the thermotaxis pathway, one of the best characterised sensorimotor circuits in *C. elegans* [16]. Previous behavioural, ablation and imaging studies have established that the thermosensory neurons AFD and AWC act through the first-layer interneurons AIY and AIZ, which converge onto the RIA [16, 17, 18]. Other classes were shown to be implicated under particular conditions. Specifically, the amphid neuron ASI contributes to thermosensory behaviour [19], AIA mediates a feeding-state-dependent pathway [20], AIB takes part in the avoidance of rapid cooling [21], RIM supports a sustained motor state [22], and the head motor neurons RMD and SMBD act down-stream of RIA [23, 18]. On the basis of these studies, we grouped the 12 classes into four layers, from sensory neurons to steering motor neurons (Fig. 1), and used this grouping only to order the classes along the pathway.

**Figure 1:**
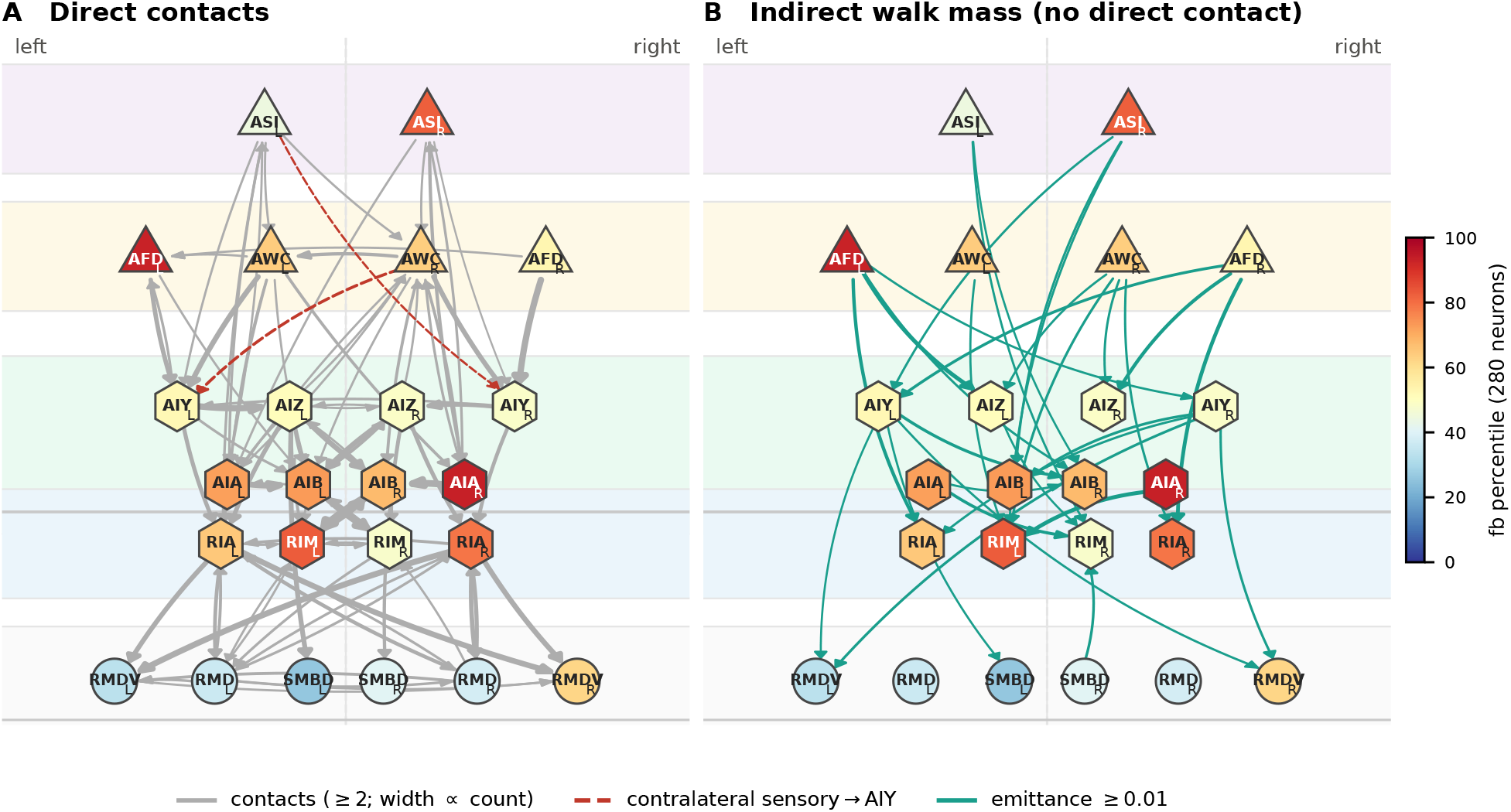
The thermotaxis and head-steering circuit. Layers from top to bottom: ASI, primary thermosensors (AFD, AWC), first-layer interneurons (AIY, AIZ, AIA, AIB), integration and motor-state interneurons (RIA, RIM), and steering output (RMD, RMDV, SMBD). Left cells on the left. Node colour: percentile of **fb** among all 280 neurons at *β*_*f*_ = 1.05 *β*_*c*_. **A**, Direct contacts (chemical synapses plus gap junctions) with at least two contacts in the merged connectome; width proportional to contact number. All contacts from AFD, AWC, and ASI onto AIY are shown; contralateral ones dashed red. **B**, Pairs without a direct contact whose emittance is at least 0.01; width proportional to emittance.

Our analysis shows that the connectome contains about three times more recurrence than strength-matched random networks, and that the excess is concentrated in sensory and weakly connected neurons, whereas random wiring places it on highly connected ones. Along the thermo-taxis pathway, recurrence is highest at the sensory end and decreases towards the integrating interneurons, while the steering classes are less consistent across reconstructions. This organisation is present in all five reconstructions we examined, including the two independently reconstructed adults, and degree-dependent loop counts do not detect it. Convergence follows the opposite pattern across the network, as the neurons with the largest excess recurrence receive their remaining input from the fewest sources. In the signal propagation atlas [7], sensory neurons show a weak bias towards incoming responses, in the same direction as the structural result, but this bias follows the number of contacts more closely than excess recurrence, so that the atlas does not resolve the recurrence gradient at the level of single neurons. Taken together, these results place the densest recurrence of the *C. elegans* connectome, relative to convergence, at its sensory periphery, a candidate structural substrate for the departure from feed-forward signalling observed in whole-brain optogenetic mapping.

## 1 Results

### 1.1 A walk-based measure of recurrence relative to convergence

Let *A* be the adjacency matrix of the directed multigraph underlying the merged *C. elegans* somatic connectome (Methods), with *N* = 280 neurons and 12,071 contact records. The entry *A*_*uv*_ is the number of chemical synapses and gap junctions from neuron *v* to neuron *u*. Let *r*(*A*) be its spectral radius and *β*_*c*_ = log *r*(*A*). For *β > β*_*c*_, the resolvent 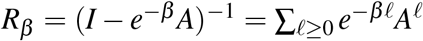 gives the weighted number of directed walks between every pair of neurons. We restrict the walk sum to walks of one or more steps, *R*_*β*_ *−I*, and define the receptance profile of neuron *u* as row *u* of *R*_*β*_ *−I* normalised to one. Its self-entry is the feedback coefficient

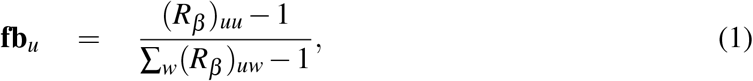

the share of the walk mass reaching *u* that started at *u*. The numerator counts the weighted closed walks that start and end at *u*, and the denominator all weighted walks that end at *u*, so **fb** expresses recurrence relative to convergence. All results are for the merged connectome at *β*_*f*_ = 1.05 *β*_*c*_ unless stated otherwise; the choice of *β*_*f*_ is explained in Methods (Fig. S2).

The feedback coefficient was only weakly related to Katz in-centrality (*ρ* = *−*0.23; Fig. 2A) and to in-strength (*ρ* = 0.11), and it followed local recurrence: the reciprocated share of a neuron’s input (*ρ* = 0.48; Fig. 2B), closed walks of length two (*ρ* = 0.51), and autapses (*ρ* = 0.51). Removing autapses changed the ranking little (*ρ* = 0.91 between the rankings with and without them). Across the connectome, the median **fb** was 0.0091. In 1,000 networks that preserved every neuron’s numbers of incoming and outgoing contacts (in- and out-strength), it was 0.0027 *±* 0.0001, and in 1,000 networks that also preserved each neuron’s number of reciprocated contacts it was 0.0041 *±* 0.0001 (*p ≤* 0.001 for both; Fig. 2C). The same comparison in each of the five reconstructions gave a median **fb** between 2.2 and 3.7 times that of its strength-preserving null networks (Fig. 2D).

**Figure 2:**
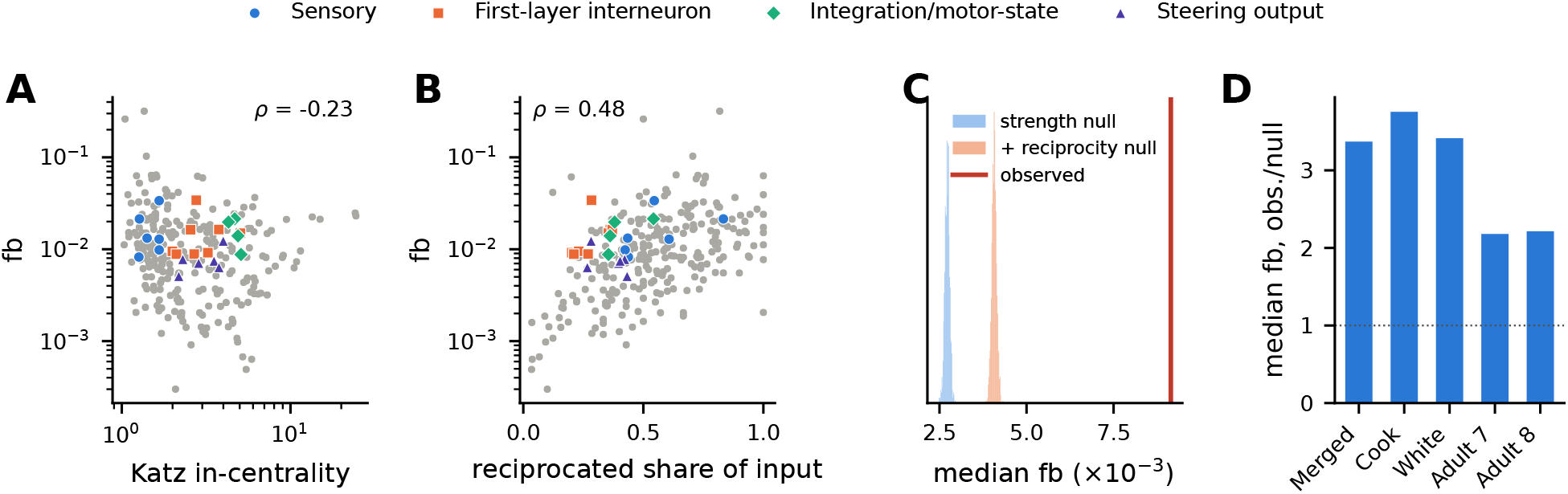
The feedback coefficient measures recurrence and exceeds strength-matched networks. **A, fb** against Katz in-centrality for all 280 neurons; circuit cells coloured by layer. **B, fb** against the reciprocated share of each neuron’s incoming contacts. **C**, Median **fb** over all neurons (red line) and in 1,000 strength-preserving (blue) and 1,000 strength- and reciprocity-preserving (orange) null networks. **D**, Ratio of observed to strength-null median **fb** in five reconstructions.

### 1.2 Excess recurrence is concentrated in sensory and weakly connected neurons

To locate the excess, we divided each neuron’s **fb** by its mean over the strength-preserving null networks. The observed value exceeded the null mean for 95% of neurons (Fig. 3A), but the excess was not uniform. It was largest in sensory neurons (median 4.4-fold; 80 neurons), smaller in interneurons (2.8-fold; 89) and motor neurons (2.6-fold; 111; sensory versus interneurons, Mann– Whitney *p* = 8 *×* 10^*−*9^), and it decreased with each neuron’s total number of contacts (*ρ* = *−*0.40, *p* = 3 *×* 10^*−*12^). The 10% most connected neurons exceeded the null 2.3-fold, the others 3.3-fold. In the null networks, **fb** increased strongly with the number of contacts (*ρ* = 0.66), so that highly connected neurons carried the most closed walks relative to their input; in the connectome this relation was weaker (*ρ* = 0.36), and weakly connected sensory neurons carried the largest excess.

**Figure 3:**
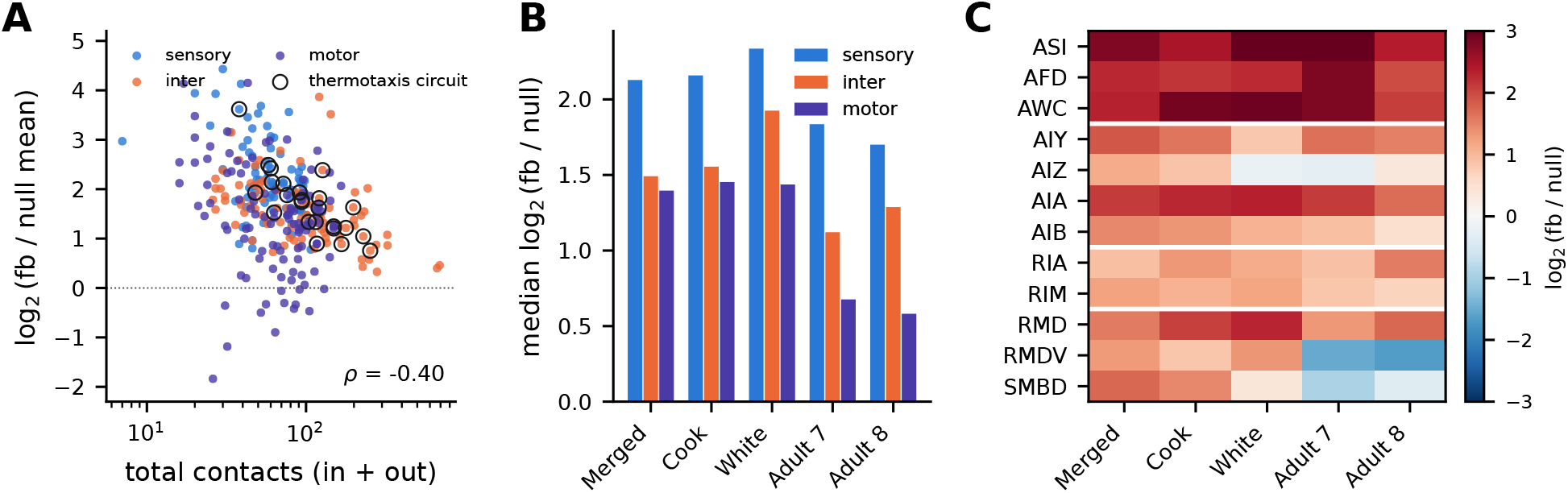
Where the excess recurrence sits. **A**, Excess recurrence (log_2_ of observed **fb** over its mean in 1,000 strength-preserving null networks) against the total number of contacts (logarithmic scale) for all 280 neurons, coloured by neuron type; thermotaxis circuit cells are circled. **B**, Median excess recurrence (log_2_) by neuron type in five reconstructions (200 null networks each for the last four). The merged, Cook, and White reconstructions are re-analyses of the original electron micrographs of White *et al*.; adults 7 and 8 are independent animals. **C**, Class-mean excess recurrence (log_2_, colour scale) for the 12 thermotaxis-circuit classes in each reconstruction; white lines separate the layers.

When each neuron’s number of reciprocated contacts was also preserved, the excess was smaller (sensory 3.0-fold, interneurons 1.9-fold, motor neurons 1.8-fold), but the difference between sensory neurons and interneurons remained (*p* = 7 *×* 10^*−*10^), while the relation with the number of contacts disappeared (*ρ* = *−*0.05, *p* = 0.43). Reciprocal contacts therefore account for the dependence on the number of contacts, and the sensory excess involves loops beyond reciprocal pairs.

The pattern was reproduced in all five reconstructions (Fig. 3B). The median excess of sensory neurons was 3.3-to 5.1-fold and that of interneurons 2.2-to 3.8-fold (sensory versus interneurons, *p ≤* 0.02 in each), and the excess decreased with the number of contacts in each reconstruction (*ρ* between *−*0.24 and *−*0.49). The strongest replication comes from adults 7 and 8 of Witvliet *et al*. [4], which are different animals reconstructed from separate electron micrographs, whereas the merged, Cook, and White reconstructions are re-analyses of the original electron micrographs of White *et al*. [1]. The pattern was also present in the network of chemical synapses alone, which excludes the bidirectional contacts contributed by gap junctions (median sensory excess 3.6-fold, interneurons 2.1-fold, *p* = 5 *×* 10^*−*9^).

### 1.3 Recurrence decreases from sensory to integrating neurons along the thermotaxis pathway

Within the thermotaxis circuit, excess recurrence decreased from the sensory layer to the integration layer, with a less consistent pattern in the downstream steering classes (Fig. 3C; Table 1). In the merged network, the class means were 8.0-, 4.9-, and 5.0-fold for ASI, AFD, and AWC; 3.6-, 4.4-, 2.7-, and 2.2-fold for AIY, AIA, AIB, and AIZ; 1.9- and 2.3-fold for RIA and RIM; and 2.5-to 3.3-fold for the steering neurons. The class-level ordering was similar across reconstructions (Spearman correlations between reconstructions 0.65 to 0.93). ASI, AFD, and AWC ranked among the five most recurrent of the 12 classes in every reconstruction, and AIZ, AIB, and RIM in the lower half in every one, while RIA was in the lower half in four of the five. The steering classes RMDV and SMBD were the least consistent: their excess was 1.3-to 1.7-fold in the three reconstructions based on the micrographs of White *et al*. and below the null in both adults. Because the adult datasets cover only the nerve ring, the contacts these neurons make outside it are missing there, which may explain the difference.

**Table 1:** Recurrent embedding of the 24 thermotaxis-circuit cells at *β*_*f*_ = 1.05 *β*_*c*_ (merged connectome). Cells are grouped by layer. In-str.: number of incoming contacts (chemical synapses and gap junctions), excluding autapses. **fb**: feedback coefficient, in units of 10^*−*3^. Pct.: percentile of **fb** among the 280 neurons. Null 95%: 2.5th–97.5th percentiles of that percentile across 1,000 strength-preserving null networks. Excess (str.) and Excess (recip.): observed **fb** divided by its mean across the strength-preserving and the reciprocity-preserving null networks, respectively. **fb**, no aut.: **fb** recomputed after removing autapses from the network (10^*−*3^).

| Cell | Layer | In-str. | **fb** | Pct. | Null 95% | Excess (str.) | Excess (recip.) | **fb**, no aut. |
|---|---|---|---|---|---|---|---|---|
| ASIL | Sensory | 16 | 8.2 | 44 | [21, 57] | 3.8 | 2.6 | 8.3 |
| ASIR | Sensory | 12 | 21.4 | 82 | [15, 48] | 12.3 | 6.0 | 21.4 |
| AFDL | Sensory | 33 | 33.8 | 93 | [81, 89] | 4.4 | 3.8 | 17.2 |
| AFDR | Sensory | 33 | 9.8 | 53 | [18, 48] | 5.4 | 3.5 | 9.9 |
| AWCL | Sensory | 23 | 13.2 | 64 | [26, 61] | 5.6 | 4.0 | 13.2 |
| AWCR | Sensory | 28 | 12.9 | 63 | [37, 69] | 4.3 | 3.0 | 12.9 |
| AIYL | First layer | 52 | 9.5 | 52 | [36, 67] | 3.3 | 2.9 | 9.5 |
| AIYR | First layer | 57 | 8.8 | 48 | [27, 59] | 3.8 | 3.2 | 8.9 |
| AIZL | First layer | 94 | 9.2 | 51 | [68, 81] | 1.9 | 1.8 | 9.2 |
| AIZR | First layer | 63 | 8.9 | 49 | [48, 73] | 2.5 | 2.2 | 8.9 |
| AIAL | First layer | 81 | 16.3 | 72 | [41, 70] | 5.2 | 4.1 | 16.4 |
| AIAR | First layer | 81 | 34.1 | 93 | [86, 92] | 3.5 | 3.2 | 14.7 |
| AIBL | First layer | 102 | 16.4 | 73 | [80, 86] | 2.3 | 2.1 | 12.3 |
| AIBR | First layer | 127 | 15.0 | 67 | [67, 81] | 3.1 | 2.7 | 14.9 |
| RIAL | Integration | 130 | 14.0 | 65 | [83, 89] | 1.7 | 1.6 | 14.0 |
| RIAR | Integration | 113 | 19.7 | 79 | [86, 91] | 2.1 | 1.9 | 16.3 |
| RIML | Integration | 83 | 21.4 | 83 | [85, 91] | 2.4 | 2.1 | 15.1 |
| RIMR | Integration | 93 | 8.8 | 48 | [53, 74] | 2.3 | 1.9 | 8.8 |
| RMDL | Steering | 63 | 7.1 | 36 | [35, 65] | 2.5 | 1.9 | 7.1 |
| RMDR | Steering | 62 | 7.4 | 38 | [24, 56] | 3.4 | 2.4 | 7.4 |
| RMDVL | Steering | 90 | 6.3 | 33 | [21, 51] | 3.1 | 2.4 | 6.4 |
| RMDVR | Steering | 85 | 12.3 | 62 | [79, 85] | 1.9 | 1.6 | 4.7 |
| SMBDL | Steering | 37 | 5.1 | 26 | [17, 46] | 2.9 | 1.8 | 5.1 |
| SMBDR | Steering | 45 | 7.8 | 41 | [22, 54] | 3.7 | 2.5 | 7.8 |

We summarised this ordering as the Spearman correlation between layer (1 to 4) and class-mean **fb** across the 12 classes. The observed value was *ρ* = *−*0.46 in the merged network, whereas the strength-preserving null predicted no ordering (0.06 *±* 0.08), as did the reciprocity-preserving null (0.00 *±* 0.08; *p ≤* 0.001 for both; Fig. 4B). In each of the other reconstructions, the observed correlation was between *−*0.24 and *−*0.44 and the null mean between 0.05 and 0.40 (*p ≤* 0.005). In the network of chemical synapses alone, the correlation was *−*0.59 against a null of *−*0.05 *±* 0.08. The five classes of the core thermotaxis pathway (AFD, AWC, AIY, AIZ, and RIA) are too few for a formal test on their own, but their observed correlations were below the null means in all five reconstructions.

**Figure 4:**
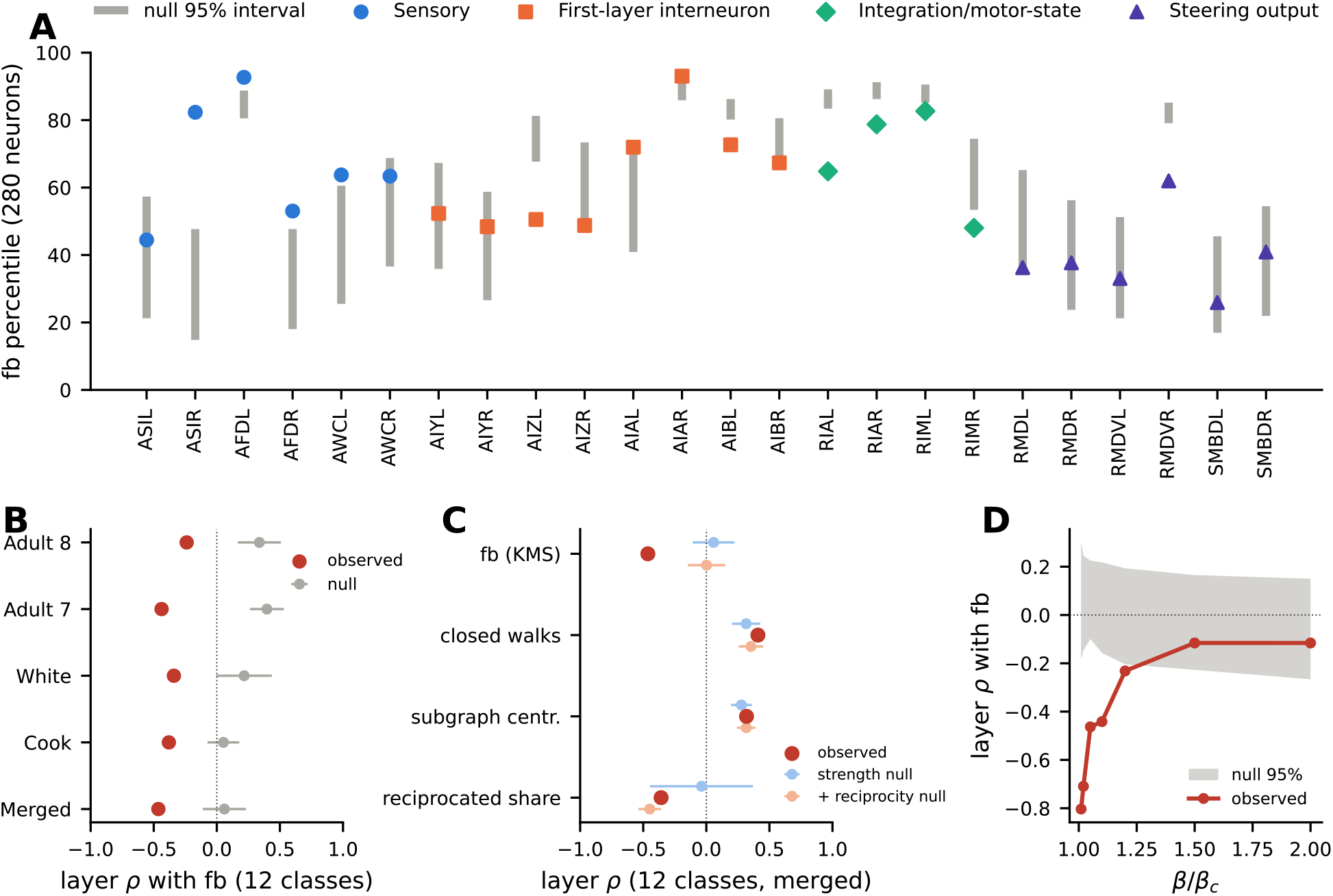
Recurrence is highest at the sensory end of the thermotaxis pathway. **A**, Percentile of **fb** among the 280 neurons for each thermotaxis circuit cell, coloured by layer, with the 2.5th–97.5th percentile range of the same percentile across 1,000 strength-preserving null networks (grey bars). **B**, Spearman correlation between layer (1 to 4) and class-mean **fb** across the 12 circuit classes in each of the five reconstructions (red), with null means *±*1.96 SD (grey). **C**, The same correlation in the merged network for **fb** and for three loop measures that are not normalised by incoming walk mass (closed walks, subgraph centrality, and reciprocated share of input), compared with the strength-preserving (blue) and reciprocity-preserving (orange) nulls (means *±*1.96 SD). **D**, The correlation for **fb** as a function of *β/β*_*c*_ (red), with the mean *±*1.96 SD of 200 strength-preserving null networks at each scale (grey band).

At the level of single cells (Fig. 4A), AFDL, AFDR, AWCL, ASIR, and both AIA cells had **fb** percentiles above the 95% interval of their percentiles across null networks, and RIAL, RIAR, RIML, RIMR, AIZL, AIBL, and RMDVR had percentiles below it. Cell-level agreement between reconstructions was moderate for **fb** itself (Spearman 0.78 between the two adults), in part because autapses, which contribute to **fb**, are recorded 77 times in the merged network but only 14 to 18 times in the White network and the adults. The class-level excess was more consistent (0.69 to 0.93 between the adults and the other networks).

### 1.4 What the normalisation contributes

Three measures of loop content that do not divide by incoming walk mass showed no ordering beyond that of the null networks (Fig. 4C). The weighted number of closed walks through each neuron, (*R*_*β*_ )_*uu*_ *−* 1, and subgraph centrality [24] both increased from sensory to integration layers (*ρ* = 0.41 and 0.32), as the null networks predicted (0.32 *±* 0.06 and 0.28 *±* 0.04; *p* = 0.13 and 0.20), because they mainly reflect each neuron’s number of contacts. The reciprocated share of a neuron’s input did decrease across layers (*ρ* = *−*0.36), but the reciprocity-preserving null reproduced this decrease (*−*0.45 *±* 0.05), and since the **fb** gradient persisted under that same null, reciprocal contacts alone do not account for it. The ordering therefore depends on comparing a neuron’s loops with the total walk mass it receives, which is what the receptance normalisation provides.

The ordering also depended on the path scale (Fig. 4D). It was strongest close to the critical value (*ρ* = *−*0.80 at 1.01 *β*_*c*_, against a null of 0.06 *±* 0.12), exceeded the null at every scale up to 1.2 *β*_*c*_, and was no longer distinguishable from the null at 1.5 *β*_*c*_ and above, where the walk sum is dominated by paths of one or two steps. Near *β*_*c*_, the denominator of **fb** is dominated by long walks from the densely connected core of the network, which we expect to reach integrating interneurons in large amounts and sensory neurons in small amounts. The ordering would then reflect the balance between local loops and global convergence, and it becomes visible only when walks of several steps are included.

The loops themselves were short (Fig. 5A). Two-step cycles, in which a neuron and one partner contact each other, accounted for 61% to 71% of the closed-walk mass in each layer, against 16% to 17% in strength-preserving networks and 26% to 43% in reciprocity-preserving networks. Long walks thus matter mainly for the denominator of **fb**, the convergent input, while the recurrence in the numerator is carried by short loops. The loops of sensory neurons were closed mainly through first-layer interneurons and other sensory neurons (Table S2). For ASIL, 35% of its closed-walk mass passed through AIAL; for AFDR, 34% through AIYR; for AWCL, 26% through AIYL; and for AWCR, 20% through AIAR. The loops of RIA passed mostly through head motor neurons (SMDV), and those of RIM through AIB.

**Figure 5:**
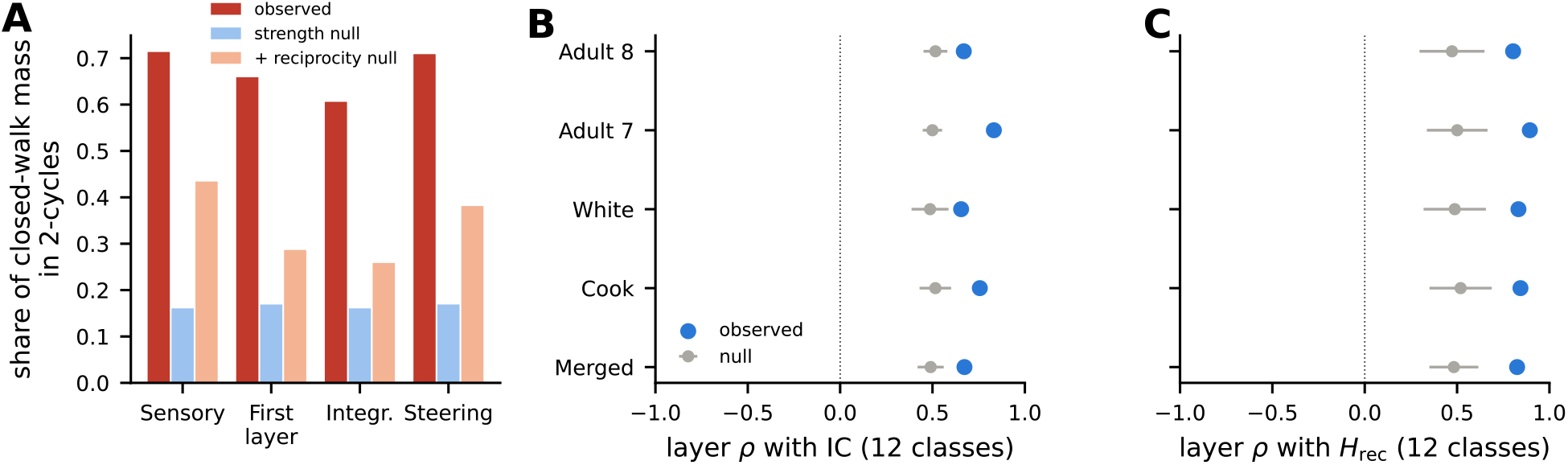
Loop composition and the complementary convergence gradient. **A**, Mean share of the closed-walk mass carried by two-step cycles for the thermotaxis-circuit cells of each layer, in the merged network (red) and in strength-preserving (blue) and reciprocity-preserving (orange) null networks. **B, C**, Spearman correlation between layer (1 to 4) and the class mean of integration capacity IC (B) or receptance entropy *H*_rec_ (C) across the 12 circuit classes in each of the five reconstructions (blue), with the mean *±*1.96 SD of the same correlation in strength-preserving null networks (grey; 1,000 networks for the merged connectome, 200 for the others).

### 1.5 Convergence is narrowest where recurrence is highest

The complement of recurrence is convergence from many sources, which we describe with two quantities. The emittance profile [13] of a source neuron *v* is column *v* of *R*_*β*_ *− I* normalised to one,

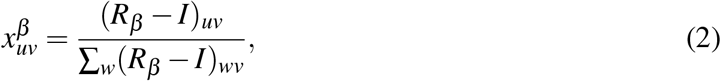

the share of the walk mass leaving *v* that reaches *u*. The integration capacity of neuron *u* is the mean share it receives from all other neurons,

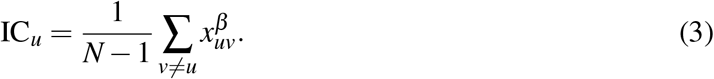

The receptance entropy is the Shannon entropy of the receptance profile of *u* with its self-entry removed and the remaining entries renormalised,

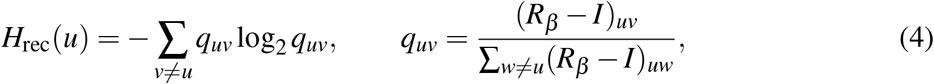

and measures how many sources contribute to the walk mass that *u* receives. Because *H*_rec_ excludes the self-entry, it is not tied to **fb** by construction.

Across the connectome, convergence showed the reverse pattern of recurrence (Fig. 6). The receptance entropy was lower than in strength-preserving networks for every one of the 280 neurons, so the walks reaching a neuron come from fewer sources than in random wiring with the same numbers of contacts. The deficit was about twice as large in sensory neurons (median 1.49 bits) as in interneurons (0.73 bits) and motor neurons (0.61 bits; sensory versus interneurons, *p* = 2 *×* 10^*−*7^), and it was unrelated to the number of contacts (*ρ* = 0.00). Sensory neurons also received less emittance than in the null networks (median IC 0.62 times the null mean), interneurons somewhat less (0.77 times), and motor neurons slightly more (1.11 times; sensory versus interneurons, *p* = 0.003). Both differences were nearly unchanged when reciprocated contacts were also preserved (*H*_rec_ deficits 1.50, 0.72, and 0.61 bits; IC ratios 0.60, 0.75, and 1.17; *p ≤* 5 *×* 10^*−*4^).

**Figure 6:**
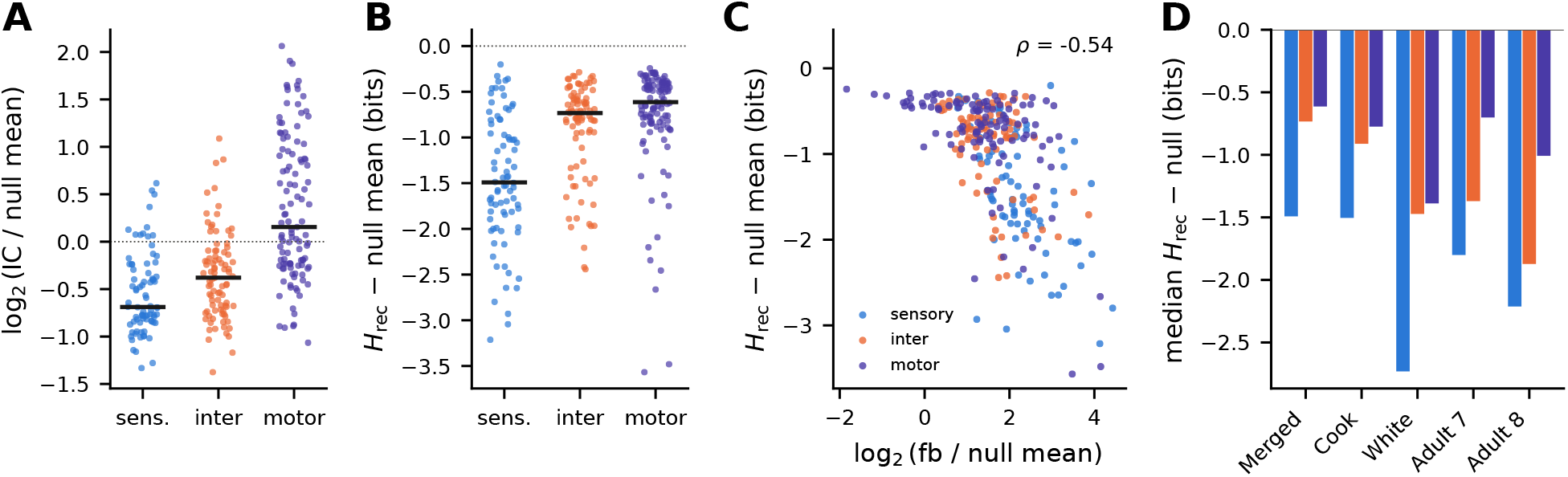
Convergence across the connectome mirrors recurrence. **A**, Integration capacity relative to its mean in 1,000 strength-preserving null networks (log_2_ ratio) for all 280 neurons, grouped by neuron type; horizontal bars are medians. **B**, Receptance entropy minus its mean in the same null networks (bits), grouped as in A. **C**, Deviation of receptance entropy from the null against excess recurrence (log_2_ of **fb** over its null mean), coloured by neuron type; *ρ*, Spearman correlation. **D**, Median deviation of receptance entropy from the null by neuron type in five reconstructions (1,000 null networks for the merged connectome, 200 for the others); colours as in C.

Recurrence and convergence were also linked neuron by neuron. Across the 280 neurons, excess recurrence was negatively correlated with the deviation of *H*_rec_ from the null (*ρ* = *−*0.54; *−*0.59 with the number of contacts controlled for; *−*0.79 under the reciprocity-preserving null) and with the IC ratio (*ρ* = *−*0.36; *−*0.45 with the number of contacts controlled for; Fig. 6C). Neurons with the most excess recurrence thus tend to receive the rest of their input from the fewest sources.

The pattern replicated in the four other reconstructions (Fig. 6D). In each of them, *H*_rec_ was below the null in at least 99.7% of neurons, the deficit was larger in sensory neurons than in interneurons (*p ≤* 0.04 in three; *p* = 0.15 in adult 8), IC relative to the null was lower in sensory neurons than in interneurons (*p ≤* 0.01), and the correlation between excess recurrence and the *H*_rec_ deviation was between *−*0.40 and *−*0.54.

Along the thermotaxis pathway, both quantities increased from sensory neurons to integration interneurons more strongly than in the null networks (IC: *ρ* = 0.67 against 0.49 *±* 0.04; *H*_rec_: *ρ* = 0.83 against 0.49 *±* 0.07; *p ≤* 0.001), and they did so in each of the five reconstructions (*p ≤* 0.005; Fig. 5B,C). IC was closely related to Katz in-centrality (*ρ* = 0.96), and *H*_rec_ less so (*ρ* = 0.75). Together with the recurrence pattern, these results describe a network in which sensory neurons receive walk mass from comparatively few sources, with a larger share than expected returning through loops they take part in, whereas interneurons and motor neurons receive it from many sources that do not return to them.

### 1.6 The signal propagation atlas does not resolve the structural pattern at single-neuron scale

We compared the structural result with the wild-type signal propagation atlas of Randi *et al*. [7]. The atlas covers the head: 186 neurons of the somatic connectome were measured, in 18,205 ordered pairs, of which 927 (5%) responded at *q <* 0.05. AWC is excluded because the atlas indexes it by ON/OFF identity, and pharyngeal neurons because they are not part of the somatic connectome. A response in the atlas reflects propagation over all paths between the stimulated and the responding neuron. For each of the 145 neurons measured in at least 20 pairs as a target and 20 pairs as a source, we computed the fraction of measured partners whose stimulation evoked a response in it (incoming response density, *d*_in_), the fraction of measured partners that responded to its stimulation (outgoing response density, *d*_out_), and the functional in/out balance (*d*_in_ *−d*_out_)*/*(*d*_in_ + *d*_out_).

Sensory neurons had a higher incoming than outgoing response density, in the direction reported by Dvali *et al*. [5] (median balance 0.09; 32 of 57 neurons; one-sided paired Wilcoxon test, *p* = 0.04, *q* = 0.06), whereas interneurons and motor neurons were balanced (medians *−*0.01 and 0.01; Fig. S3A). The bias was of similar size at response thresholds of *q <* 0.01 and *q <* 0.1 (*p* = 0.06 and 0.05), and the balance did not differ significantly between sensory and other neurons (Mann–Whitney *p* = 0.23). Across neurons, the balance was weakly correlated with excess recurrence (*ρ* = 0.15, *p* = 0.08) and more strongly, and negatively, with the number of contacts (*ρ* = *−*0.24, *p* = 0.005). With the number of contacts controlled for, it was unrelated to excess recurrence (partial *ρ* = 0.04, *p* = 0.61; Fig. S3B). Functional reciprocity, the fraction of a neuron’s responding partners that also responded in the reverse direction, was weakly and not significantly correlated with **fb** (*ρ* = 0.16, *p* = 0.08, 123 neurons; Fig. S3C), to the same extent as with the structural reciprocated share (*ρ* = 0.16) and in-strength (*ρ* = 0.18), and not at all after controlling for these (partial *ρ* = 0.02). Within the thermotaxis circuit, responses from one layer to the next and in the reverse direction had similar amplitudes (*p ≥* 0.78 for each pair of adjacent layers; Fig. S3D).

The atlas therefore does not provide single-neuron validation of the structural result. Sensory neurons show an input bias in the same direction, but it is not significant after correction for multiple comparisons and is not larger than in other neuron types. The atlas records responses to brief stimulation of one neuron at a time, only 5% of measured pairs respond, and the in/out balance depends more on the number of contacts than on excess recurrence.

## 2 Discussion

Across the *C. elegans* connectome, recurrent embedding measured relative to convergence is highest at the sensory periphery and lowest in the highly connected integrating interneurons, whereas random wiring with the same numbers of contacts places the most recurrence on highly connected neurons. Convergence follows the complementary pattern, since sensory neurons draw their incoming walks from particularly few sources and the two deviations from random wiring are correlated across the whole network. In the thermotaxis pathway this organisation appears as a gradient from ASI, AFD, and AWC to RIA and RIM, while the downstream steering classes are less consistent, and the gradient is present in all five reconstructions, including the two independently reconstructed adults.

The pattern does not follow from the simplest statistics of the wiring. It exceeds networks that preserve each neuron’s numbers of incoming, outgoing, and reciprocated contacts, it persists in the network of chemical synapses alone and therefore does not arise from the symmetry of gap junctions, and it holds for sensory neurons across the whole connectome, of which the thermotaxis circuit is one instance. The null models leave other properties of the network unconstrained, however, including the spatial arrangement of neurons and the rich-club organisation of its hubs [6], and part of the excess may reflect such constraints. Contacts were also treated as unsigned and chemical synapses and gap junctions were weighted equally, so the analysis says where returning walks are concentrated but not whether the corresponding loops are excitatory or inhibitory, which would decide whether they amplify or attenuate sensory activity.

The result depends on how recurrence is measured. Closed walk counts and subgraph centrality, which increase with the number of contacts in the same way as in the null networks, do not detect it, and it appears only when a neuron’s closed walks are divided by all the walk mass it receives. The underlying matrix is the Katz resolvent, and the difference lies in this normalisation, which Katz centrality and communicability do not perform because they sum incoming walks without separating the part that returns to the neuron. The gradient also depends on the path scale, being strongest close to the critical value and indistinguishable from the null when only walks of one or two steps contribute. Since most of the closed walk mass is carried by two-step loops, we interpret the role of long walks as acting through the denominator of **fb**, as walks from the densely connected core of the network add to the convergent input of integrating interneurons more than to that of sensory neurons. This interpretation is consistent with the convergence results but was not tested directly, and the gradient exceeds the null only for 1.01 *≤ β/β*_*c*_ *≤* 1.2.

Some aspects of the replication are weaker than others. The class-level ordering agreed well between reconstructions, whereas cell-level values agreed only moderately, in part because autapses, which contribute to **fb**, are recorded with very different frequencies in different reconstructions. We therefore reported **fb** with and without autapses. The adult datasets cover only the nerve ring, so neurons with processes outside it lose part of their connectivity, which may explain why the steering classes differed between the adults and the other reconstructions. The assignment of the 12 thermotaxis classes to four layers follows the literature and orders the classes along the pathway, but the whole-network result, which compares neuron types and does not use the layers, does not depend on it.

The loops of the thermosensory neurons pass mostly through their first-layer targets, AFD through AIY, AWC through AIY and AIA, and ASI through AIA, with further loops among sensory neurons such as ASE and ASG. Reciprocal connections from AIY to AFD and AWC have been proposed to shape thermosensory responses [17], and AIA integrates sensory and state information from several amphid neurons [20]. The structural result suggests that such feedback is a general property of sensory neurons in this network and extends beyond the individual pairs studied so far, whereas integrating interneurons receive convergent input from many neurons that do not receive their output in return, which is consistent with a role in pooling signals.

These findings bear on the divergence between the connectome and the signalling network reported by Dvali *et al*. [5], whose observations that signalling is not feed-forward along the sensory-to-motor axis and that amphid sensory neurons receive more functional connections than they send suggest that the sensory periphery is where signalling departs most from feed-forward flow. The structural analysis places the largest excess of returning walks, relative to what a neuron receives, at the same periphery. In the signal propagation atlas, sensory neurons showed a weak input bias in the same direction, but the bias was not significant after correction for multiple comparisons and was not larger than in other neurons, the in/out balance followed the number of contacts more than excess recurrence, and functional reciprocity followed structural reciprocity. The structural and functional observations therefore do not yet connect at the level of single neurons. Extrasynaptic signalling, which the connectome does not contain, contributes to the atlas [7], and responses to brief stimulation of single neurons may not engage loops that take several steps to close. Whether sensory recurrence shapes ongoing sensory dynamics, adaptation, or responses to repeated stimulation will require other experiments, for example comparing AFD or AWC responses to temperature steps with and without AIY or AIA activity, and signed connectivity estimates would indicate whether the recurrent pathways identified here are predominantly excitatory or inhibitory.

## 3 Methods

### 3.1 Connectomes

All analyses were first carried out on the merged somatic connectome of the hermaphrodite from [14], in which the network of Varshney et al. [2] is completed with 3,900 connections from Cook et al. [3]. It contains 280 neurons and 12,071 contact records of unit weight, 77 of them autapses, with every gap junction entered in both directions. The chemical-only and gap-junction-only networks were extracted from the typed version of the same file. For replication we used the somatic network of Cook et al. [3], together with the reconstruction of White et al. [1] and the adult datasets 7 and 8 of Witvliet et al. [4] in the form distributed with WormNeuroAtlas (v0.0.7.3). The Witvliet datasets cover the nerve ring only, and gap junctions that a source lists once were inserted in both directions. Neuron types (sensory, interneuron, motor) follow the annotation distributed with the KMSnet code, and the thermotaxis classes and circuit layers are those described in the Results.

### 3.2 Walk-based quantities

For an adjacency matrix *A* with spectral radius *r*(*A*), the resolvent *R*_*β*_ = (*I − e*^*−β*^ *A*)^*−*1^ is finite for every *β > β*_*c*_ = log *r*(*A*), which equals 4.296 for the merged network. It was obtained by direct matrix inversion at *β*_*f*_ = 1.05 *β*_*c*_ unless another scale is stated. The feedback coefficient **fb**, the integration capacity IC and the receptance entropy *H*_rec_ were computed from *R*_*β*_ *− I* as defined in the Results, so every profile counts walks of one or more steps. The identity is removed because the walk of length zero puts the same unit mass on the diagonal for every neuron and carries no information about the wiring. The share of closed walks of length *ℓ* in the recurrence of *u* was taken as *e*^*−βℓ*^(*A*^*ℓ*^)_*uu*_*/* ((*R*_*β*_ )_*uu*_ *−* 1 for *ℓ* = 1, The loop partners of *u* were ranked by *e*^*−β*^ *A*_*vu*_(*R*_*β*_ )_*uv*_, the closed-walk mass of *u* whose first step leads to *v*. Subgraph centrality [24], used as a comparison for **fb**, was computed as [exp(*A/r*(*A*))]_*uu*_.

### 3.3 Choice of the path scale

The scale *β* controls how quickly the weight of a walk decays with its length. The analysis needs a value at which walks of several steps contribute appreciably and the states of different neurons remain distinguishable. We examined several criteria over 1.001 *≤ β/β*_*c*_ *≤* 2.5 (Fig. S2), computed on the KMS states themselves, namely the columns of *R*_*β*_ normalised to sum to one.

As *β* decreased towards *β*_*c*_, the mean Shannon entropy of the emittance profiles rose monotonically, from 0.7 at 1.5 *β*_*c*_ to 7.0 bits at 1.001 *β*_*c*_ (Fig. S2A), and the mean receptance entropy was also largest at the smallest *β* examined. Close to *β*_*c*_ every state approaches the Perron vector of *A*, and the profiles become almost indistinguishable. Their mean pairwise Jensen–Shannon divergence is 0.008 bits at 1.001 *β*_*c*_, compared with 0.31 bits at 1.05 *β*_*c*_. Maximising the mean entropy would therefore push *β*_*f*_ onto *β*_*c*_, where the states no longer tell neurons apart.

Two other criteria peak just above *β*_*c*_. The first is the standard deviation across neurons of the emittance entropy, which measures how differently the states of individual neurons spread over the network. It had a single maximum at 1.06 *β*_*c*_ (1.43 bits, against 1.42 bits at 1.05 *β*_*c*_; Fig. S2A). The second is the mean fraction of each state lying outside its own neuron. This fraction fell from nearly 1 close to *β*_*c*_ to 0.07 at 1.5 *β*_*c*_, and it crossed 0.5, the point where the walks leaving a neuron and its self-weight carry equal mass, at 1.07 *β*_*c*_ (Fig. S2B). Its value at 1.05 *β*_*c*_ was 0.57. We took *β*_*f*_ = 1.05 *β*_*c*_ as a representative value within this narrow regime, and we do not treat it as a parameter fitted to biological data. At this scale the median **fb** (0.0091) was close to its broad maximum of 0.0094, which is reached between 1.08 and 1.11 *β*_*c*_ (Fig. S2C). The ordering from sensory to integrating layers exceeded the null at every scale from 1.01 to 1.2 *β*_*c*_ (Fig. 4D), so the conclusions do not depend on the exact value chosen.

### 3.4 Null networks

Strength-preserving null networks were generated by permuting the targets of all contact records other than autapses, which keeps the in-strength and out-strength of every neuron. Records that became self-contacts in the process were exchanged with random partners until none remained. Reciprocity-preserving null networks split each ordered pair into a reciprocated part, min(*A*_*uv*_, *A*_*vu*_) in each direction, and a one-way remainder. The targets of the one-way records were permuted, and the reciprocated records were rewired as undirected stubs and inserted in both directions, which preserves in-strength, out-strength and the number of reciprocated contacts of each neuron. Autapses were left in place in both models. Every null network was analysed at 1.05 times its own critical value. Excess recurrence is the observed **fb** divided by its mean across null networks. IC was compared with the null in the same way, whereas *H*_rec_ was compared through its difference in bits from the null mean. The merged network was compared with 1,000 networks of each type. The other reconstructions and the chemical-only and gap-junction-only variants were each compared with 200 strength-preserving networks. One-sided *p*-values were computed as (*k* + 1)*/*(*B* + 1), where *k* is the number of null networks at least as extreme as the observation and *B* the number of networks.

### 3.5 Class-level tests

The ordering across the circuit was summarised by the Spearman correlation between layer, coded from 1 at the sensory layer to 4 at the motor layer, and the class means of a given quantity over the 12 thermotaxis classes. The observed correlation was compared with the distribution of the same correlation across null networks. Neuron types were compared with two-sided Mann–Whitney tests on excess recurrence.

### 3.6 Signal propagation atlas

We used the wild-type atlas of Randi et al. [7] in the version distributed with WormNeuroAtlas (v0.0.7.3), without merging bilateral cells. Its matrices are indexed by responding target and stimulated source. The atlas contains 23,902 measured ordered pairs among head neurons, 18,205 of them between neurons of the somatic connectome. Ventral-cord motor neurons, which the atlas numbers without zero padding, were renamed to match the connectome labels. AWC was left out because the atlas identifies its two cells by their ON and OFF states rather than by side. A pair counted as measured when its *q*-value was defined and as responding when *q <* 0.05, and the thresholds *q <* 0.01 and *q <* 0.1 were used in sensitivity analyses. Self-responses were not used.

For a neuron *u*, the incoming response density is the fraction of measured sources whose stimulation evoked a response in *u*, and the outgoing response density is the fraction of measured targets that responded to the stimulation of *u*. The in/out balance is their difference divided by their sum. Neurons measured fewer than 20 times as target or as source, or without any responding pair, were excluded, leaving 145 neurons. Raising the threshold to 40 removed one neuron and did not change the results. In sensory neurons the incoming and outgoing densities were compared with a one-sided Wilcoxon signed-rank test, in the direction reported by Dvali et al. [5]. Neuron types were compared with Mann–Whitney tests. The balance was related to excess recurrence, **fb** and the number of contacts by Spearman correlations with permutation *p*-values from 9,999 permutations, and partial correlations were computed from the residuals of the ranks.

Functional reciprocity was assessed in neurons with at least 10 partners measured in both directions and at least 3 responding partners. Forward and backward responses between adjacent circuit layers were compared by the mean |Δ*F/F*| over all measured cell pairs. This comparison used a two-sided test against 5,000 permutations of the layer labels among the 11 classes present in the atlas.

### 3.7 Multiple comparisons

Table S1 lists all 41 tests reported in the paper, with *q*-values adjusted across this whole family by the Benjamini–Hochberg procedure [25].

## Code and data availability

The code used in this study is available in the KMSnet repository (https://github.com/elkMm/KMSnet). The connectome data are distributed with the repository and with WormNeuroAtlas, and the atlas data are those of Randi et al. [7].

## Acknowledgment

This work was supported by the Natural Sciences and Engineering Research Council of Canada through the CRC grant NC0981.

## Author contributions

E.M.M. conceived the study, developed the theoretical framework and computational tools, performed the analysis and wrote the manuscript. H.B. analysed results, wrote the manuscript and secured funding.

## Notes

### Competing Interest Statement

The authors have declared no competing interest.

